# Cytosolic fumarase acts as a metabolic fail-safe for both high and low temperature acclimation of *Arabidopsis thaliana*

**DOI:** 10.1101/2021.04.19.440416

**Authors:** Helena A. Herrmann, Pablo I. Calzadilla, Jean-Marc Schwartz, Giles N. Johnson

**Author notes:** Corresponding author: 0161 275 5750.

## Abstract

Plants acclimate their photosynthetic capacity in response to changing environmental conditions. In *Arabidopsis thaliana,* photosynthetic acclimation to cold requires the accumulation of the organic acid fumarate, catalysed by a cytosolic fumarase FUM2. However, the role of this accumulation is currently unknown. In this study, we use an integrated experimental and modelling approach to examine the role of FUM2 and fumarate across the physiological temperature range. Using physiological and biochemical analyses, we demonstrate that FUM2 is necessary for high as well as low temperature acclimation. We have adapted a reliability engineering technique, Failure Mode and Effect Analysis (FMEA), to formalize a rigorous approach for ranking metabolites according to the potential risk that they pose to the metabolic system. FMEA identifies fumarate as a low-risk metabolite, while its precursor, malate, is shown to be high-risk and liable to cause system instability. We conclude that the role of cytosolic fumarase, FUM2, is to provide a fail-safe, controlling malate concentration, maintaining system stability in a changing environment. We argue that FMEA is a technique that is not only useful in understanding plant metabolism but can also be used to study reliability in other systems and synthetic pathways.

## Introduction

Plants in natural environments are continually exposed to fluctuating conditions. In response to short term changes in the environment, plants can *regulate* enzymes to ensure systemic homeostasis is maintained. Over longer periods, plants *acclimate*, altering the concentration of different enzymes to match the prevailing circumstances (Webber, 1994; Stitt and Hurry, 2002). How plants sense their environment and how that leads to acclimation is poorly understood, however, there is growing evidence that metabolic signals play an important role on this process (Templeton and Moorhead, 2004; Krahmer et al. 2018; Herrmann et al. 2019a).

During the day, plants fix atmospheric carbon through photosynthesis. A large proportion of that carbon accumulates in the leaf during the photoperiod. Overnight it is then remobilised to support growth and cell maintenance. Under different environmental conditions, the way in which plants store these photoassimilates changes. For instance, the model plant *Arabidopsis thaliana* (hereafter Arabidopsis) stores carbon primarily as the polysaccharide starch and the organic acids fumarate and malate (Chia et al., 2000; Smith and Stitt, 2007; Zell et al., 2010; Pracharoenwattana et al., 2010). Starch turnover in the chloroplast has been studied extensively as this is the major non-structural carbohydrate in most plants (Smith and Stitt, 2007; Streb and Zeeman, 2012). Malate and fumarate are intermediates of the tricarboxylic acid cycle in the mitochondria, but in some species, they are also produced in the cytosol and accumulate diurnally in the vacuole (Kovermann et al., 2007; Fernie and Martinoia, 2009).

In the cytosol in Arabidopsis, malate can be made from or converted to oxaloacetate, citrate, isocitrate, cis-aconitate, α-ketaglutarate, glutamic acid, pyruvate and fumarate (Arnold and Nikoloski, 2014) providing carbon skeletons for various biosynthetic pathways, including amino acid synthesis (Zell et al., 2010). In addition, malate is a key metabolite in the redox shuttling between the chloroplast, mitochondrion, and cytosol; balancing the ATP/NADPH ratio and the cellular energy supply within the plant cell (reviewed by Selinski and Scheibe, 2018). The Malate/OAA shuttle has also been strongly linked with acclimation to different types of stresses in plant (Kandoi et al. 2017; Wang et al. 2016; Dinakar et al. 2016), for which subcellular concentrations needs to be precisely regulated.

Fumarate is typically produced from malate via the enzyme fumarase and, in addition to the mitochondrial fumarase isoform (FUM1), some plants species like Arabidopsis have a second fumarase located in the cytosol (FUM2). Cytosolic fumarate is primarily synthesized during the day and is converted back to malate during the night, acting as a diurnal carbon storage (Pracharoenwattana et al., 2010; Zell et al., 2010; Dyson et al., 2016; Pracharoenwattana et al., 2010). Fumarate accumulation in Arabidopsis leaves has previously been linked to growth on high nitrogen and nitrogen metabolism (Pracharoenwattana et al., 2010; Araujo et al. 2011) and its accumulation was observed to be proportional to biomass production at low temperature (Scott et al., 2014). However, the specific role of fumarate accumulation is still unclear.

Previously, we showed that diurnal fumarate accumulation increases immediately in response to cold in Arabidopsis and continues to increase until photosynthetic acclimation is achieved in these plants (Dyson et al., 2016; Herrmann et al., 2021). In contrast, knockout mutants lacking FUM2 expression, which do not accumulate fumarate (Pracharoenwattana et al., 2010), are unable to acclimate their photosynthetic capacity in response to low temperature (Dyson et al., 2016). The role of fumarate accumulation in the low temperature acclimation of Arabidopsis remains elusive.

Photosynthetic responses to temperature significantly vary between species (Way and Yamori, 2014). Generally, when a plant is transferred from warm to cold conditions, enzymatic activities are reduced, slowing metabolism. Under such conditions, light absorption can exceed the capacity for assimilation of the energy, triggering the production of reactive oxygen species (ROS), photosystem II (PSII) photoinhibition and cell damage (Murata et al. 2007; Miura and Furumoto, 2013; Ding et al. 2019). Over subsequent days, following acclimation, plants transferred to the cold typically increase the concentration of key enzymes, increasing the capacity for carbon assimilation that acts as an electron sink (Adams et al. 2013). Following acclimation, plants achieve a similar rate of photosynthesis at low temperature to that previously seen in the warm (Hurry et al., 2000; Strand et al., 2003; Dyson et al., 2016). This acclimation process is reflected in their maximal photosynthetic capacity (P_max_), which is higher in cold treated plants when compared to those kept under control conditions (Athanasiou et al., 2010; Dyson et al., 2016; Herrmann et al. 2019b).

At high temperature, photosynthesis is initially limited by alterations in the PSII repair cycle and the activation state of Rubisco (reviewed by Allakhverdiev et al. 2008; Salvucci and Crafts-Brandner, 2004). Rubisco activation is regulated by Rubisco activase, which is a thermo-labile enzyme whose activity decreases when temperature increases (Jensen, 2000; Eckardt and Portis, 1997; Crafts-Brandner and Salvucci, 2000). Hence, photosynthetic acclimation to heat may imply an increase in Rubisco activase content (Liu and Huang, 2008; Wang et al. 2010). However, at sub-lethal high temperatures (up to around 30 °C), photosynthetic capacity increases with increasing temperature. Way and Yamori (2014) reported that several plant species reduce their photosynthetic capacity when growth at high temperatures, what is defined as detractive adjustment. Although these adjustments might not be beneficial from a photosynthetic perspective, it might benefit the plant performance, in terms of optimising their nitrogen allocation (Way and Yamori, 2014).

To gain a deeper insight into the role of fumarate accumulation in photosynthetic acclimation to temperature, we have here analysed the photosynthetic and biochemical responses of three Arabidopsis genotypes that accumulate fumarate to varying degrees. Comparisons across the physiological temperature range (5-30 °C) were carried out on Col-0 wild type, C24 wild type (which has a reduced capacity of fumarate accumulation) and the *fum2* mutant. We demonstrate that FUM2 activity is essential not only for low temperature acclimation but also for high temperature responses. To further understand the role of fumarate accumulation, we have adapted a reliability engineering technique, Failure Mode and Effect Analysis (FMEA, Stamatis, 1995; Rausanda and Øienb, 1996), to formalize a rigorous approach for ranking metabolites according to the potential risk that they pose to the metabolic system. We employ both a small-scale metabolic model and a genome-scale pathway analysis to conclude that cytosolic fumarase FUM2 acts as a fail-safe, maintaining system stability under changing environmental conditions. We argue that our FMEA adaptation is not only useful in understanding plant metabolism but can also be used to study reliability in other metabolic systems.

## Materials and Methods

### Plant growth and tissue preparation

*A. thaliana* Col-0, C24 and *fum2.2* genotypes were grown in peat-based compost in 3-inch pots with 8-hour photoperiods at 20 °C day/18 °C night, and under a 100 μmol m s irradiance, as previously described (Dyson et al., 2016). Light was provided by warm white LEDs (colour temperature 2800-3200 K). Adult plants (8-9 weeks) were then either kept at 20/18 (day/night) °C for up to another week (control treatment) or transferred to 5/5, 10/8, 15/13, 25/23, or 30/28 (day/night) °C, under identical irradiance and photoperiod conditions. Fully expanded leaves were collected at the beginning and at the end of the photoperiod on the 1^st^ and 7^th^ day of the different treatments, flash-frozen in liquid nitrogen, freeze-dried and stored at −20 °C for subsequent metabolite assays. Additionally, leaves were also collected without freezing and their fresh and dry weights per unit area recorded.

### Gas exchange

Photosynthesis and respiration were measured under growth conditions (ambient temperature and light intensity) in plant growth cabinets, using a CIRAS 1 infra-red gas analyser fitted with a standard broad leaf chamber (PP systems, Amesbury, USA), as previously described (Dyson et al., 2016). Measurements were taken on the 1^st^ and 7^th^ day for the cold (5 °C) and warm (30 °C) treatment, and under control conditions. The maximum capacity for photosynthesis (P_max_) was estimated at 20 °C and under 2000 ppm CO_2_ and 2000 µmol m^-2^ s^-1^ light, as described previously (Herrmann et al., 2019b). All measurements were taken between the 5^th^ and 7^th^ hour of the photoperiod and on a minimum of 4 biological replicates.

### Metabolite assays

Fumarate, malate and starch concentrations were estimated as previously described by Dyson et al. (2016). Averages and standard errors of 3-4 replicates were calculated, and an analysis of variance (ANOVA) followed by a Tukey’s test was performed in R, with a confidence level of 0.95. Where measurements were subtracted from one another, the uncertainty (μ) was calculated such that 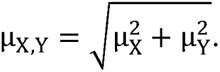 Measurements where the uncertainty ranges do not overlap are considered to be significantly different.

### Kinetic modelling

We used kinetic modelling to assess how changes in metabolic fluxes correspond to changes in metabolite concentrations. Kinetic modelling is not practical with large networks. Thus, rather than using a genome-scale model we constructed a minimal model based on a sub-set of reactions (Fig. 1; Table S1). For simplicity, we did not consider compartmentalization. Kinetic reactions were set up in COPASI (Version 4.27.217). Rates of photosynthesis and respiration were set as independent variables for Col-0, C24 and *fum2* plants, after being converted to mmol CO_2_ (gDW)^-1^ s^-1^ (Fig. S1). We initially parametrized the model to fit the measured average diurnal carbon fluxes to starch, malate, and fumarate for control conditions, using simple mass action kinetics (Abegg, 1899). All other metabolites were assumed not to accumulate in the leaf and were constrained to concentrations between 0-0.0005 μmol CO_2_ (gDW) s. The rate of carbon export was allowed to adjust freely, accounting for any remaining carbon.

**Figure 1:**
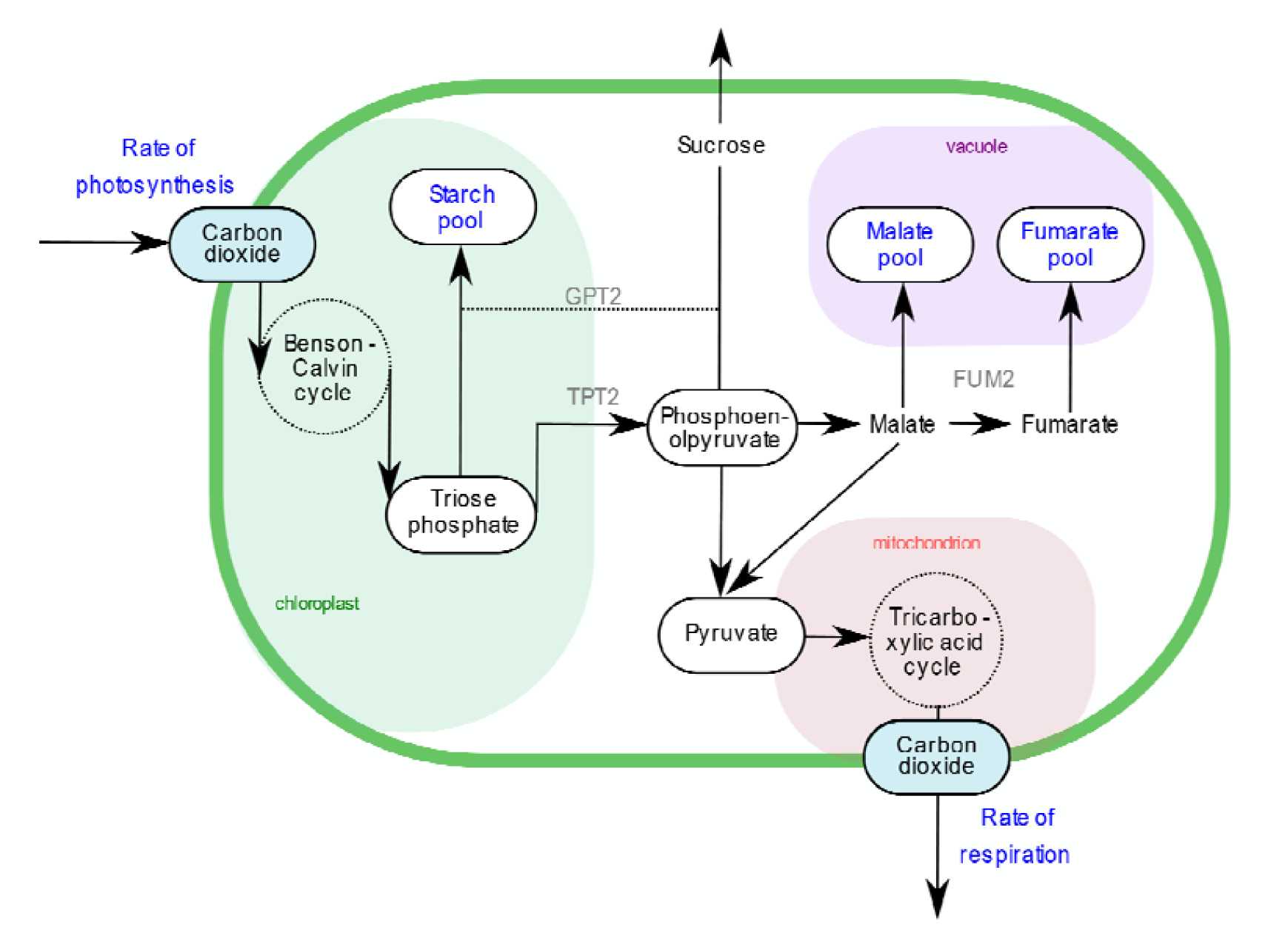
Illustration of the reactions used to set up the kinetic model. Metabolite concentrations, shown as ovals, and reactions, shown as black solid lines, were considered in the model as described (see Materials and Methods). Metabolite pools and the rates of photosynthesis and respiration (as shown in blue) were experimentally determined for each genotype and used to parametrize the kinetic parameters in the model fitting. For simplicity, compartmentalization was not considered in the kinetic model and the diurnal sugar accumulation was assumed to be minimal, as shown previously (Dyson et al. 2016).

Using the Hooke and Jeeves (1961) parameter estimation algorithm, with an iteration limit of 10^3^, a tolerance of 10^-8^ and a rho of 0.2, we found a model solution for which the estimated concentration values fell within the uncertainty ranges of the experimentally measured values. We used an Arrhenius constant to capture temperature-dependence and allowed the effective Q_10_ to vary between 1.0-3.0 to fit the model to the measured starch, malate, and fumarate concentrations at T = 5 °C and T = 30 °C. Without implementing any regulatory mechanisms, we were able to find a solution for which the concentration values estimated by the model fell within the uncertainty ranges of the experimentally measured values (Fig. S2). It is important to note that the effective Q_10_, as represented in our model, quantifies a possible temperature sensitivity of a reaction rather than the intrinsic properties of the enzymes.

### Failure Mode and Effect Analysis (FMEA)

An existing genome-scale metabolic model of Arabidopsis plant leaf metabolism (Arnold and Nikoloski, 2014; Herrmann et al. 2019b) was converted to a directed metabolite-metabolite graph and analysed using the networkx package (Version 2.2) in Python (Version 3.6.9). Reversible reactions were accounted for by applying two separate edges between the respective metabolites, one in either direction. All non-carbon compounds, small molecules and cofactors were removed from the graph. The model is leaf specific and is compartmentalized to include the chloroplast, mitochondrion, cytosol and peroxisomes.

We applied the FMEA framework to primary carbon metabolism. We considered the metabolites as the components of the metabolic system. We calculated the risk of failure for each component, considering failure to be that the concentration of a particular metabolite falls outside a concentration range required to sustain metabolic activity. The FMEA analysis does not presume what the required concentration of a metabolite should be at a given condition and can therefore be considered an unbiased approach (Lewis et al., 2012).

The probability of failure for a given metabolite M downstream of ribulose-1,5-bisphosphate carboxylase/oxygenase (Rubisco) was calculated as:

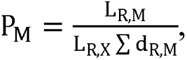

where L_R,M_ is equal to the shortest path length from Rubisco to metabolite M, L_R,X_ is the longest of all the shortest paths to metabolites in the network, and ∑ d_R,M_ is the number of alternative paths to metabolite M having the shortest length. L_R,X_ is a normalization constant and is equal to 14 in all calculation of P_M_. Overall, P_M_ considers the probability of failure of M as a calculation of the number of upstream reactions and metabolites that are required for the production of M.

The severity of failure for each metabolite (S_M_) was calculated such that:

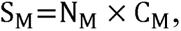

where N_M_ is the degree of metabolite M (i.e. number of direct neighbours) and C_M_ is the normalized betweenness centrality of M. The number of neighbours of M indicates the number of reactions that would fail if M was not present in the system. Betweenness centrality is defined as the normalized sum of the total number of shortest paths that pass through a node and can thus be considered a proxy for how important that metabolite is in the production of other metabolites. S_M_ calculates a severity for M based of the number of reactions and down-stream metabolites that cannot occur in the absence of M.

The risk factor (R_M_) was calculated for each metabolite such that

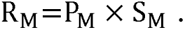

The risk factor takes into consideration how heavily the production of M is dependent on other metabolites and other reactions in the system. It also considers how many other metabolites and reactions in the system are dependent on the production of M. Therefore, R_M_ considers how important M is to the overall system functionality.

Although we could equally well calculate the risk factor of individual reactions in a metabolic system, we have here chosen to consider metabolites as the individual components of the system. This is because metabolites are the fundamental building blocks of the metabolic system, whereas reactions are the means to producing these building blocks. For example, it is possible for one reaction to fail but for all metabolites in the system to still be produced as required.

### Data availability

All experimental data, model parameterizations and network analyses are available on Github (https://github.com/HAHerrmann/FMEA_KineticModel) and under the following Zenodo DOI: 10.5281/zenodo.3596623

## Results & Discussion

Photosynthetic acclimation involves different mechanisms and processes, which play a role in the adjustment of plant metabolism to a changing environment (Herrmann et al. 2019a). In terms of temperature, photosynthesis shows different responses when plants are exposed to heat or cold, although some of their signalling pathways are shared (reviewed by Sharma et al. 2020; Choudjury et al. 2013). Previously, we observed that photosynthetic acclimation to cold is dependent on the presence of FUM2 activity or protein (Dyson et al., 2016; Herrmann et al. 2020). Thus, with the aim of understanding further the role of FUM2 and fumarate in the photosynthetic acclimation response to temperature, we analysed how its accumulation changes across the physiological temperature range in different genotypes of Arabidopsis.

### Leaf diurnal fumarate accumulation increases in response to both low and high temperature

Malate and fumarate represent important diurnal carbon stores in Arabidopsis, and their accumulation tends to increase at low temperature (Dyson et al. 2016; Herrmann et al. 2019b; Herrmann et al. 2020). The Arabidopsis ecotype C24 has previously been reported to have a reduced capacity for cytosolic fumarate synthesis, while the knockout mutant *fum2* is impaired in fumarate accumulation (Riewe et al., 2016; Pracharoenwattana et al. 2010). To understand the impact of the accumulation of these molecules across the physiological temperature range, we examined beginning and end of day leaf concentrations of malate and fumarate in Col-0, C24 and the *fum2* plants.

Diurnal accumulation was estimated in plants grown at 20 °C and in plants transferred to a range of temperature, on the 1^st^ (Fig. 2) and the 7^th^ (Fig. 3) day of treatment. Under controlled conditions, both malate and fumarate accumulate in an approximately linear fashion across the 8-hour photoperiod (Dyson et al., 2016). In Col-0 on the 1^st^ day of treatment, malate accumulation was unaffected by temperature (Fig 2a), however fumarate accumulated to significantly higher concentrations at low, but also at high temperatures (Fig. 2d). In C24, malate accumulation was similar to that in Col-0 (Fig. 2b), while fumarate accumulation was lower at all temperatures, with this increase only differing significantly from the control at the lowest temperature measured. Nonetheless, an increasing trend of fumarate accumulation at both low and high temperature was observed (Fig. 2e). There was no significant change in fumarate concentration across the photoperiod in *fum2*, however malate accumulation did vary significantly between some temperatures, again showing an increasing trend both at low and high temperatures (Fig. 2c,f).

**Figure 2:**
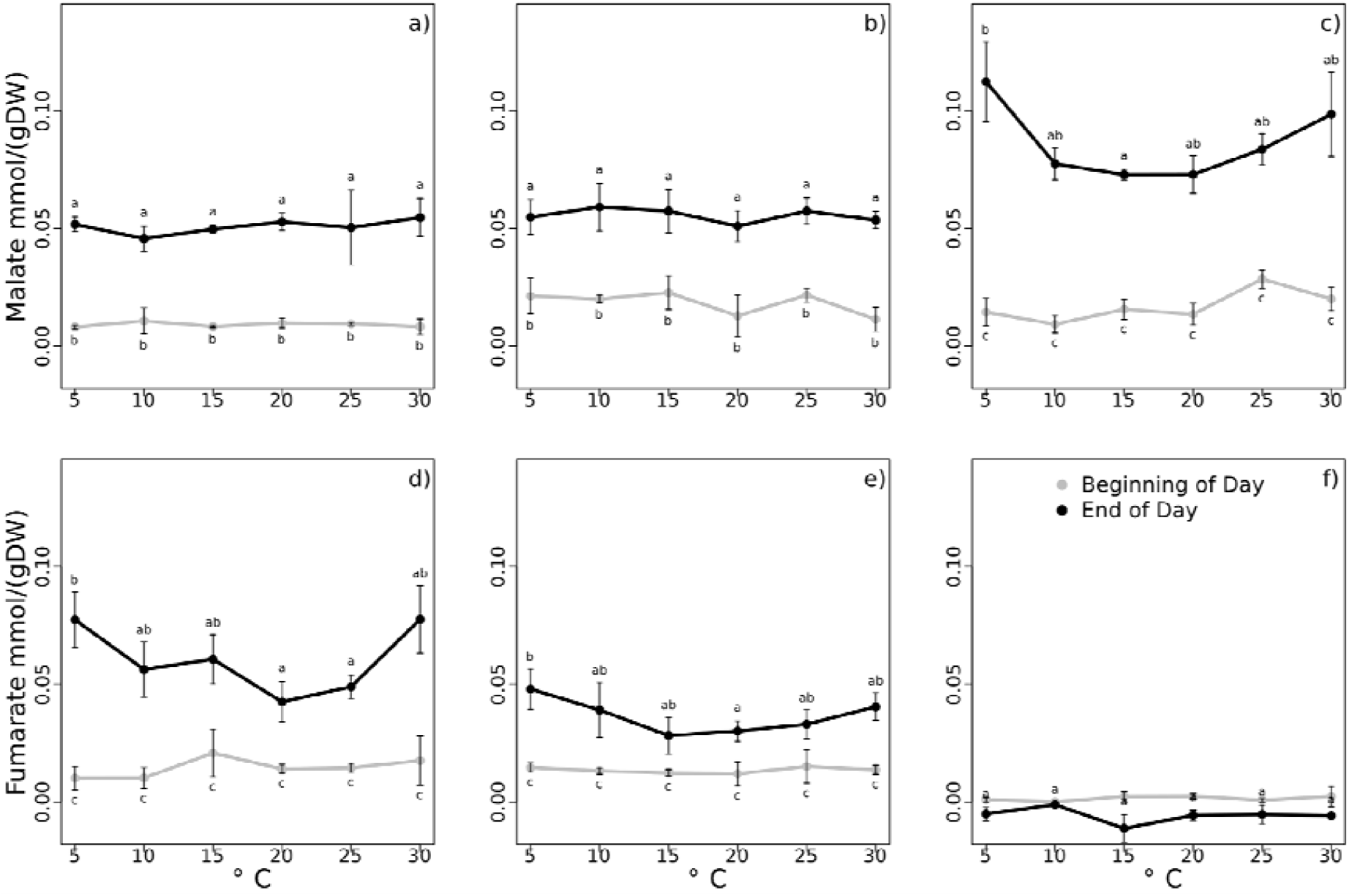
Organic acid concentrations of three Arabidopsis genotypes measured across 6 temperatures on the 1^st^ day of treatment. Beginning of day (BOD; grey) and end of day (EOD; black) concentration of malate and fumarate are shown for Col-0 (a,d), C24 (b,e) and *fum2* (c,f) genotypes. Standard mean errors of 3-4 biological replicates are shown. Different letters above the error bars indicate statistically different values (Analysis of variance, Tukey’s test with a confidence level of 0.95).

**Figure 3:**
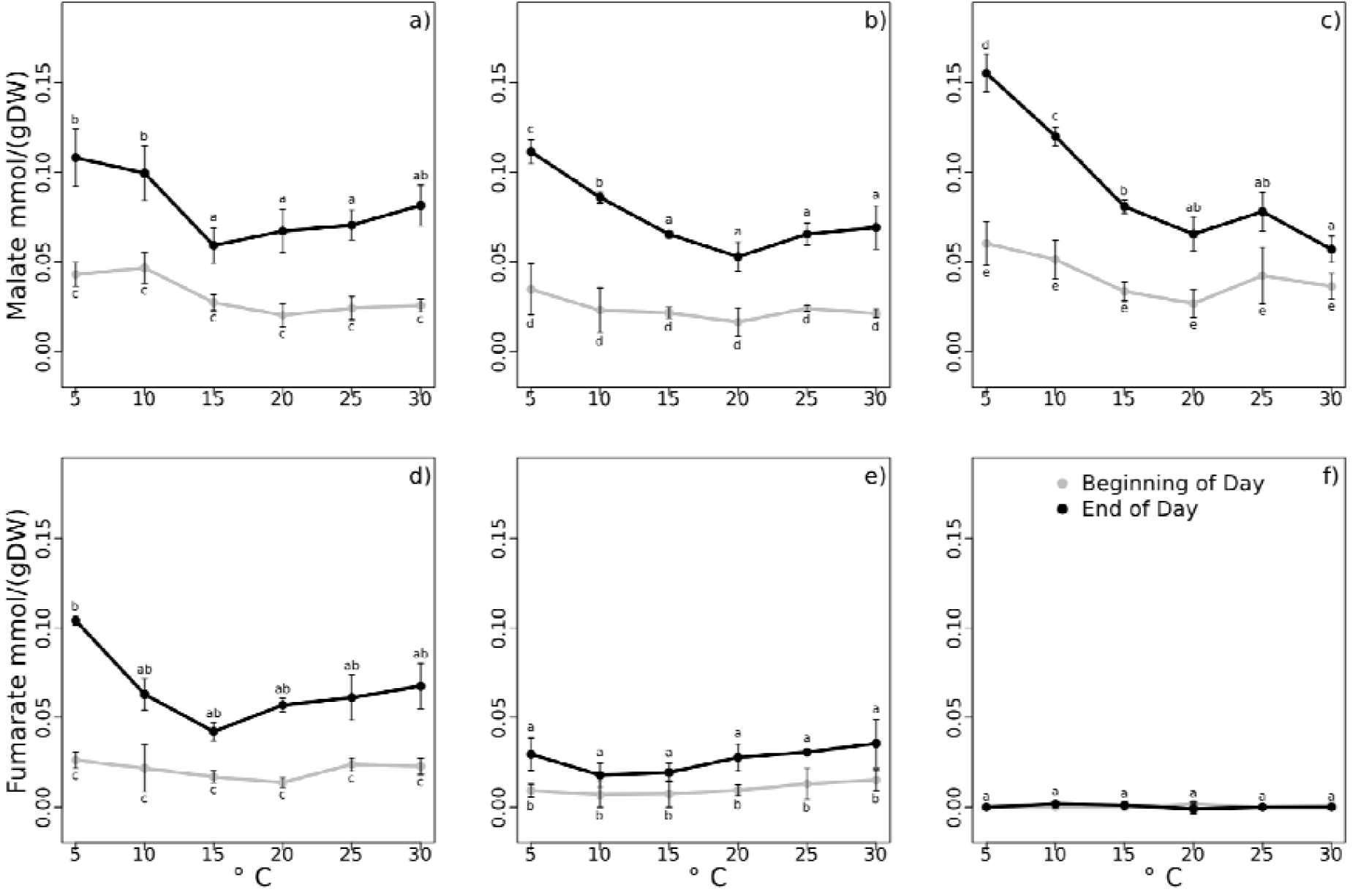
Organic acid concentrations of three Arabidopsis genotypes measured across 6 temperatures on the 7^th^ day of treatment. Beginning of day (BOD; grey) and end of day (EOD; black) concentration of malate and fumarate are shown for Col-0 (a,d), C24 (b,e) and *fum2* (c,f) genotypes. Standard mean errors of 3-4 biological replicates are shown. Different letters above the error bars indicate statistically different values (Analysis of variance, Tukey’s test with a confidence level of 0.95).

Following 7 days of cold treatment, malate accumulation was higher than in control conditions in all plants, whilst under warm treatment, malate concentration remained constant (Fig. 3a-c). Fumarate accumulation remained high in Col-0 after 7-days of cold treatment but fell back to control levels in C24 (Fig. 3d,e). After 7 days of high temperature treatment, fumarate accumulation was not significantly different to control conditions (Fig. 3d-f).

The third major diurnal carbon store in Arabidopsis is starch (Smith and Stitt, 2007; Streb and Zeeman, 2012). We previously demonstrated that the total accumulation of malate, fumarate and starch in Col-0 and *fum2* changes upon cold acclimation (Dyson et al. 2016, Herrmann et al. 2019c, Herrmann et al. 2020). We measured starch accumulation at the lowest and highest temperatures and observe the same trend, with the addition of the C24 accession (Supplementary Figure S3). Combining data from Fig’s 2,3 and S3, we estimated the total leaf diurnal carbon storage, which also increases its total diurnal carbon accumulation upon cold acclimation (Fig. 4a). Upon heat treatment, however, we did not observe a change in the this (Fig. 4b).

**Figure 4:**
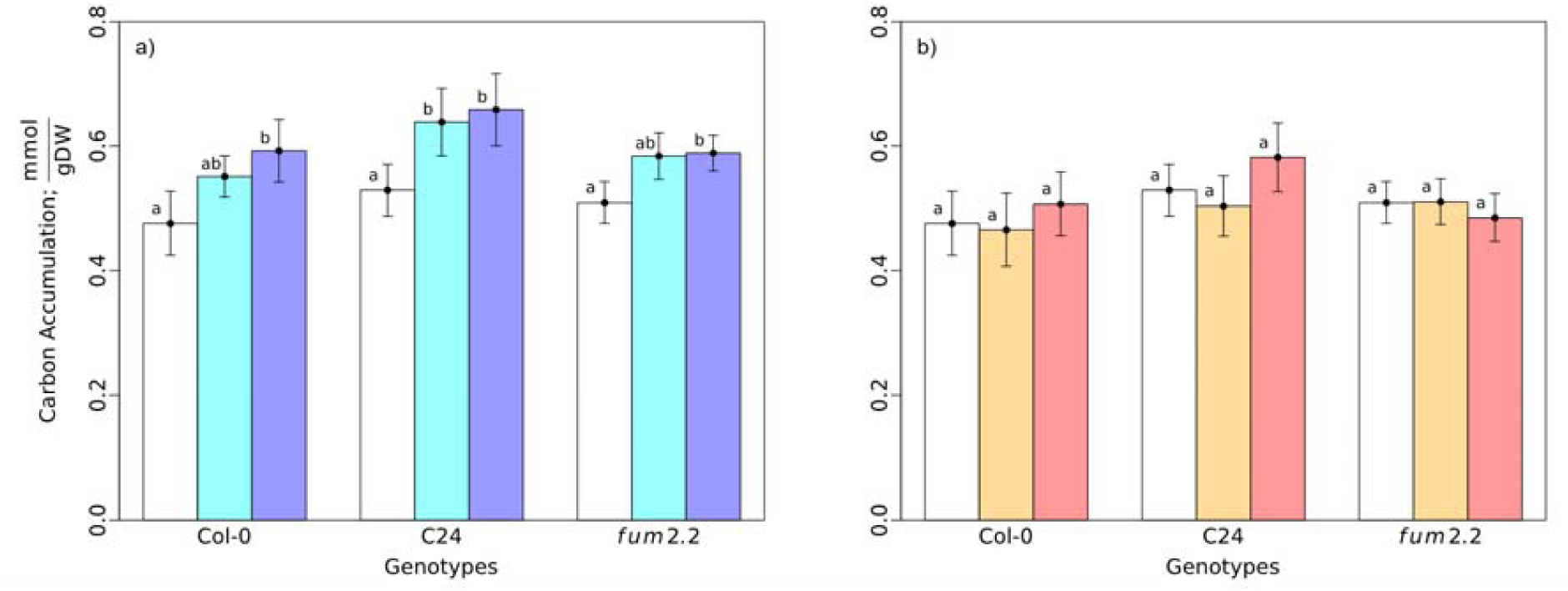
Total diurnal carbon accumulation, calculated from the combined accumulation of malate, fumarate and starch (Fig. S3), in Col-0, C24, and *fum2* plants under control conditions (white) or after one day (cyan or orange) and seven days (blue or red) of cold (a) and warm (b) treatments, respectively. Diurnal accumulation was calculated by subtracting the beginning of day concentrations from the end of day concentrations. The calculated uncertainties are shown as mean error bars and were used to check for significance (see Materials and Methods for further details).

### Photosynthetic acclimation to both low and high temperatures requires FUM2 activity

When exposed to a change in their environment, plants adjust their maximum photosynthetic capacity (P_max_) in order to optimize metabolism to the new prevailing condition. P_max_ (measured under common control conditions) provides an indication of the acclimation state of the photosynthetic apparatus, giving the maximum photosynthetic capacity of plants at saturating CO_2_ and light conditions (Herrmann et al. 2019b). Previously, we demonstrated that *fum2* plants are unable to acclimate their photosynthetic capacity to cold, establishing a link between fumarate accumulation via FUM2 and photosynthetic acclimation (Dyson et al. 2016; Herrmann et al. 2019c). To test whether changes in fumarate accumulation also impair photosynthetic acclimation under other temperature conditions, we examined the photosynthetic capacity of the three tested Arabidopsis accessions across the full physiological temperature range (Fig. 5). Col-0, C24, and *fum2* were grown at 20 °C for 8 weeks, and either kept at 20 °C or transferred to 5, 10, 15, 25 or 30 °C for 7 days. P_max_ was measured at 20 °C and under light- and CO_2_-saturating conditions on the 1^st^ (Fig. S4) and 7^th^ (Fig. 5) day of treatment.

**Figure 5:**
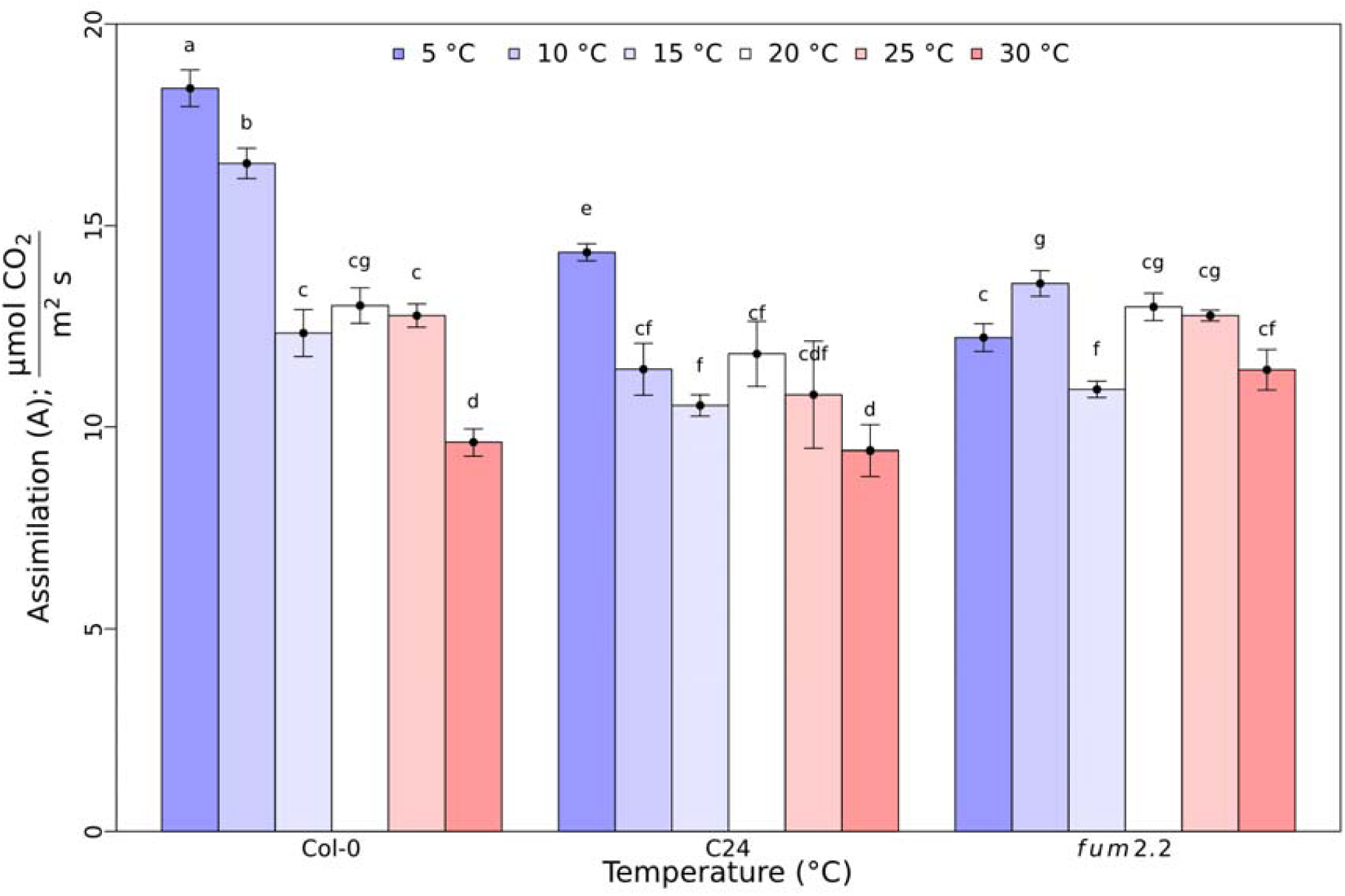
Maximum capacity for CO_2_ assimilation (*P_max_*) was measured at 20 °C and under light- and CO_2_-saturating conditions on adult plants grown and at 20 °C and then exposed to 5, 10, 15, 20 25, or 30 °C treatments for one week. Different letters above the error bars indicate statistically different values (Analysis of variance, Tukey’s test with a confidence level of 0.95).

P_max_ did not change on the 1^st^ day of treatments nor did it vary significantly between genotypes (Fig. S1). However, after one week of treatment, Col-0 and C24 show a significant change in P_max_ in response to the 5, 10, and 30 °C treatments (Fig. 5). In both genotypes, P_max_ increases in response to cold and decreases in response to warm treatment, albeit not to the same extent. The *fum2* mutant, which lacks cytosolic fumarase, does not show a consistent trend in altering P_max_ in response to temperature (Fig. 5).

P_max_ measurements provide information about the relative investment of the plant in the photosynthetic apparatus, but they do not reveal the actual rates of photosynthesis in growth conditions. Thus, CO_2_ fixation under growth conditions was evaluated on the 1^st^ and 7^th^ day of cold (5 °C) and warm (30 °C) treatments (Fig. 6). In plants transferred to cold, CO_2_ fixation was initially inhibited in all three genotypes. In Col-0 and C24 plants, photosynthesis recovered after one week of acclimation, reaching the same rate as observed in control conditions (Fig. 6a). In contrast, the *fum2* mutant failed to recover its rate of photosynthesis in the cold. Under warm treatment (Fig. 6b), Col-0 plants showed a lower rate of photosynthesis after one week, whereas C24 and *fum2* plants maintained a constant rate of photosynthesis throughout the treatment.

**Figure 6:**
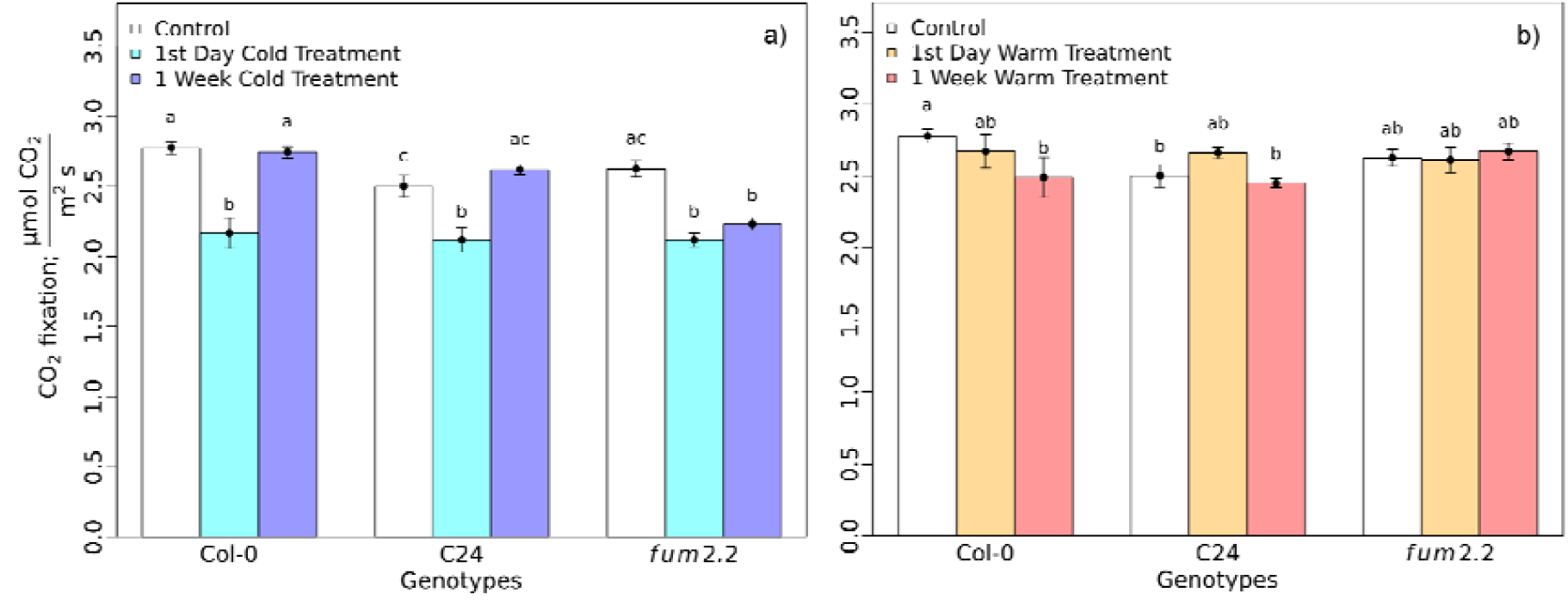
In-cabinet rates of photosynthesis measured on the 1^st^ and 7^th^ day of the cold (a) and warm treatments (b). CO_2_ fixation measurements were taken under ambient light in adult Arabidopsis plants grown under control temperature conditions (white), after one or seven days of cold treatment (cyan and blue) and warm treatment (orange and red), respectively. Standard mean errors of 3-4 biological replicates are shown. Different letters above the error bars indicate statistically different values (Analysis of variance, Tukey’s test with a confidence level of 0.95).

Our results show that diurnal fumarate accumulation increases in response to both cold and warm treatment in wild-type plants (Fig. 2&3), and this is necessary for photosynthetic acclimation under both conditions (Figure 5). C24 plants accumulate intermediate lower of fumarate than the Col-0 wild-type and show an attenuated temperature acclimation of P_max_. Meanwhile, *fum2* plants are unable to accumulate fumarate and do not show a consistent adjustment of P_max_ in response to either low or high temperatures.

### The role of fumarate in the temperature acclimation of photosynthesis

Fumarate accumulation could either act as a signal for acclimation itself or could modulate metabolic or redox responses to temperature (Dyson et al. 2016; Herrmann et al. 2020). If the first hypothesis were correct, we would expect similar changes in fumarate concentration to trigger similar outcomes for photosynthetic acclimation. However, the increased fumarate accumulation at high and low temperatures are accompanied by opposite photosynthetic acclimation responses. As such, we can exclude a direct signalling role for fumarate itself.

If fumarate is not directly sensing temperature to regulate acclimation, we must consider how fumarate accumulation indirectly modulates other signals. Together, the organic acids fumarate and malate form an important diurnal carbon sink for photosynthates. For instance, eliminating the flux to fumarate (*fum2* knockout genotype) results in an increase in the accumulation of malate at both low and high temperature during the first day of treatment (Figure 4a & b). Plants with reduced fumarase activity (C24), however, show no significant change in malate across the temperature range (Fig. 2) though they do accumulate more starch (Fig. S3e-f). Total diurnal C storage in the leaf is similar in all three genotypes under any given condition but this is distributed differently between storage pools.

Increasing the metabolic fluxes to different carbon sinks, such as starch, is a relevant strategy to avoid photoinhibition under stress conditions, like low temperatures (Ensminger et al. 2006; Adams et al. 2013; Allakhverdiev et al. 2008). However, total carbon accumulation did not significantly change between the studied genotypes under the tested temperature conditions (Fig. 4), implying that C24 and *fum2* are not limited by their sink capacity. Thus, the link between fumarate accumulation and photosynthetic acclimation might not be merely related to its function as a carbon sink and instead is likely to be connected to a more complex metabolic response.

### Increased fumarate accumulation under high and low temperatures does not requires regulatory mechanisms

We have previously reviewed how mathematical modelling can be used as a powerful tool to study plant metabolism and its connection to photosynthetic acclimation (Herrmann et al. 2019a, Gjindali et al. 2021). The observation that flux to fumarate increases under both low and high temperature conditions was unexpected and shows that the flux to fumarate itself does not correlate directly to a change in temperature, as previously speculated (Dyson et al. 2016). To understand how the observed fumarate accumulation links to temperature-dependent changes in enzyme kinetics, we constructed a kinetic model of the primary reactions in leaf carbon metabolism (Fig. 1, Table S1). To keep the number of reactions in the kinetic model to a minimum, only relevant branch and end points were included in the model.

In comparison to starch, malate and fumarate, neither sucrose nor amino acids accumulate substantially in Arabidopsis leaves in our growth conditions (Dyson et al., 2016); however, we can assume that there is a significant flux through these pools that constitutes phloem export, with sucrose representing the most important flux (Lalonde et al., 2003, Ainsworth et al., 2011). Measured rates of photosynthesis and respiration (Fig. 6; Fig. S5) were used to constrain the model. The remaining parameters were estimated to fit the measured malate, fumarate and starch concentrations in control conditions and on the 1^st^ day of 5 and 30 °C treatment, respectively. It is assumed that on the 1^st^ day of temperature treatment, plants will not yet have significantly acclimated to the stress condition as no change in P_max_ is observed (Fig. S4). Thus, observed metabolic changes are largely the result of kinetic or regulatory effects rather than changes in enzyme concentrations due to acclimation. When fitting separate model instances for Col-0, C24 and *fum2* plants, a solution with the same kinetic parameters for all reactions in the three genotypes is found, except for the conversion of fumarate to malate and that of triose phosphate to starch (Table 1). The models predict no cytosolic fumarase activity in *fum2* and a reduced activity in C24 compared to Col-0 plants, consistent with experimental observations.

**Table 1.**
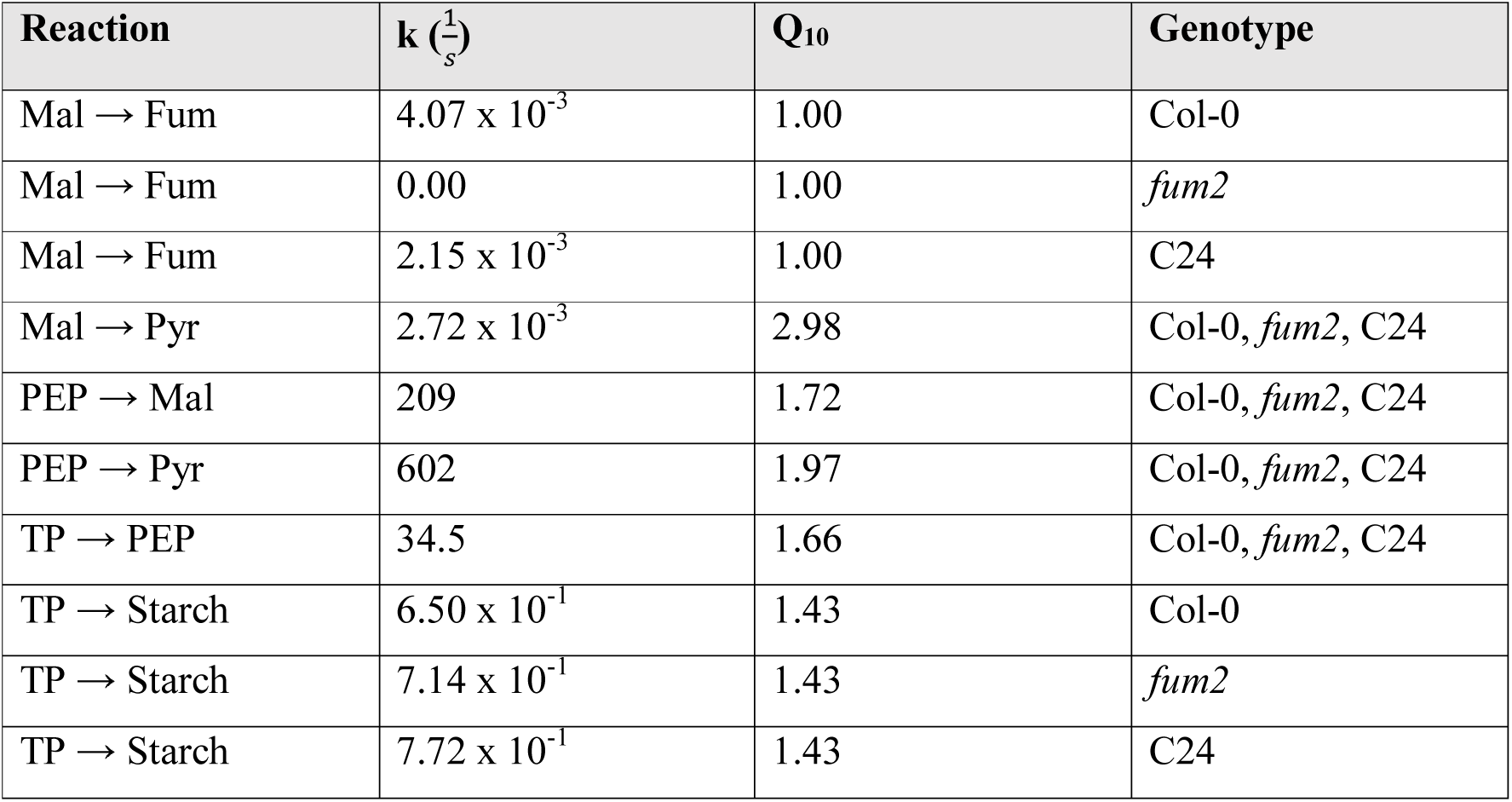
Parameters for a kinetic model of carbon metabolism. The metabolites malate (Mal), fumarate (Fum), pyruvate (Pyr), phosphoenolpyruvate (PEP), triose phosphate (TP) were used to construct the model. The full table including the rates of photosynthesis and respiration set as model constraints are shown in the Supplementary Materials (Table S1).

Most of the effective Q_10_ values are estimated at around 2.0 (Table 1), as is typical for temperature-dependence of biological reactions (Elias, 2014). However, the cytosolic fumarase reaction shows an apparent temperature independence, with an effective Q_10_ = 1.0, indicating that the corresponding enzyme is not limiting under any of the modelled conditions. The model results demonstrate that differences in the rate constants are enough to explain the observed difference of metabolite accumulation between genotypes. The increased fumarate production in response to both warm and cold treatment can result simply from temperature-dependent enzyme kinetics, without implying another kind of regulation of enzymatic activity. Although we cannot exclude that regulatory effects are occurring, the model shows that a solution for organic acid production without temperature-dependent regulatory effects is possible.

The conversion of malate to pyruvate (Fig. 1) is, according to the model solutions, the most temperature-sensitive reaction; however, it carries little overall flux. The conversions of phosphoenolpyruvate to malate and pyruvate, on the other hand, carry a substantial flux and show a strong temperature-dependence. The counter-play of these two reactions, along with the observed changes in rates of photosynthesis, respiration, and carbon export (Fig. S6), are sufficient to account for the experimentally observed changes in organic acid concentrations under warm and cold temperature treatments.

Thus, our mathematical modelling confirms that an increased accumulation of fumarate and a constant accumulation of malate are metabolically plausible under the Arrhenius law (Arrhenius, 1889), and that no regulatory mechanisms are required for fumarate accumulation to increase at both high and low temperatures. Nevertheless, active regulation of fumarate accumulation under these conditions cannot be excluded. In this regard, metabolite imports and exports from different organelles, which was not consider due to the required complexity of the modelling approach, could also play a critical role in fumarate regulation. For this reason, we also considered a pathway analysis of a genome-scale model of Arabidopsis leaf metabolism.

### Fumarate as a fail-safe for carbon metabolism homeostasis under different environmental conditions

The above results suggest a passive, rather than an active role for fumarate accumulation in the plant responses to temperature but does not tell us the role of fumarate. Using an existing metabolic model of all of Arabidopsis leaf primary carbon metabolism, we have taken a network topology approach to understand how organic acid accumulation changes with temperature treatment. Adapting a Failure Mode and Effect Analysis (FMEA; see Materials and Methods), we were able to identify metabolites that pose a high and low risk for the ‘failure’ of the metabolic system. High-risk metabolites are those that, in the event of failure, will create the greatest disturbance to the metabolic system. In engineering, a risk factor is calculated based on the probability of failure (P_M_) and the severity of failure (S_M_). Here, we estimated P_M_ based on the length of the shortest path and the number of paths that lead to a metabolite M. S_M_ can be associated with the number of neighbours of a metabolite and the number of paths that lead to a metabolite (see Materials and Methods). The latter is calculated as the betweenness centrality (i.e., the number of shortest paths that pass through that metabolite).

Table 2 shows the metabolites with the 10 highest risk factors obtained from our analysis. The average risk factor of all metabolites was 0.005. Malate, oxaloacetate and glucose-6-phosphate stand out as high-risk metabolites in the cytosol (Table 2). In the chloroplast, the metabolites with the highest risk factor were α-ketoglutarate, glyceraldehyde 3-phosphate, pyruvate, fructose 6-phosphate, ribose 5-phosphate, glucose 6-phosphate, and phosphoenolpyruvate. Mitochondrial metabolites related to carbon metabolism have an average or below average risk factor. Only the chloroplast, mitochondrion, cytosol and peroxisomes compartments were taken into consideration, as specified in the Arnold and Nikoloski (2014) model. The full list of risk factor calculations are available on Zenodo and Github (DOI: 10.5281/zenodo.3596623).

**Table 2:** The ten most high-risk metabolites as identified by the Failure Mode and Effect Analysis (FMEA). An existing metabolic model (Arnold and Nikoloski, 2014; Herrmann *et al*., 2019c) was converted to a graph of carbon metabolism as outlined in the methods. Metabolites with the highest risk factors (*R*) and their cellular locations are shown. Possible cellular locations, as written in the genome-scale metabolic model, include the chloroplast, mitochondrion, cytosol and peroxisomes. R=S*_M_* × *P_M_*, where *S_M_*=*C_M_* × *N_M_*, was calculated such that the severity is equal to the normalized number of neighbours (*N_M_*) times the normalized betweenness centrality (*C_M_*). PM is based on the length of the shortest path and the number of shortest paths that lead to a metabolite *M*.

| Metabolite | Location | $N_M$ | $C_M$ | $P_M$ | $R$ ($\times 10^{-2}$) |
| --- | --- | --- | --- | --- | --- |
| Pyruvate | Chloroplast | 0.778 | 0.206 | 0.357 | 5.71 |
| Phosphoenolpyruvate | Chloroplast | 0.389 | 0.129 | 0.286 | 5.71 |
| α-Ketoglutarate | Chloroplast | 0.944 | 0.124 | 0.357 | 4.19 |
| Glyceraldehyde 3-phosphate | Chloroplast | 0.833 | 0.134 | 0.286 | 3.24 |
| Malate | Cytosol | 1.00 | 0.0374 | 0.500 | 1.87 |
| Glucose 6-phosphate | Cytosol | 0.500 | 0.0695 | 0.500 | 1.74 |
| Ribose 5-phosphate | Chloroplast | 0.500 | 0.0976 | 0.357 | 1.74 |
| Fructose 6-phosphate | Chloroplast | 0.667 | 0.0685 | 0.357 | 1.63 |
| Oxaloacetate | Cytosol | 1.00 | 0.0564 | 0.429 | 1.42 |
| Glucose 6-phosphate | Chloroplast | 0.389 | 0.0727 | 0.429 | 1.21 |

The network structure that connects all high-risk metabolites shows that these are well-connected metabolites and are located only a few reactions upstream of a carbon sink (Table 2, Fig. 7). Carbon sinks are generally low-risk metabolites (R < 0.001). In particular, fumarate is an endpoint in the system and has a null betweenness centrality (C_M_). Therefore, changing the concentration of fumarate has a negligible effect on the overall metabolic system. Because changes in the concentration of high-risk metabolites could have a negative effect on system functionality, we hypothesized that adjusting the flux to the nearest carbon sink could minimize the overall system disturbance. Thus, for instance, fumarate could act as a fail-safe, buffering changes in malate, oxaloacetate and other upstream metabolites under different temperature conditions (Fig. 7). The observation that fumarate accumulation is necessary for photosynthetic acclimation to temperature is consistent with flux to fumarate acting as an inherent fail-safe to the metabolic system. Fumarate itself was defined as a low-risk metabolite, and changing its concentration is presumed to have a negligible effect on overall system functionality. Thus, an influx of carbon can be re-directed to fumarate without disturbing metabolism. However, malate and oxaloacetate, the immediate precursors of fumarate (Fig. 7), were identified as high-risk components and alterations in their concentrations might cause system failure. As a consequence, their accumulation levels need to be tightly regulated. Accordingly, unlike fumarate, the amount of malate accumulating through the day in both wild type accessions is surprisingly constant in response to initial warm and cold treatment and only changes following acclimation (Fig. 2&3). By contrast, the inability of *fum2* plants to accumulate fumarate results in increasing malate levels and their lack of photosynthetic acclimation capacity.

**Figure 7:**
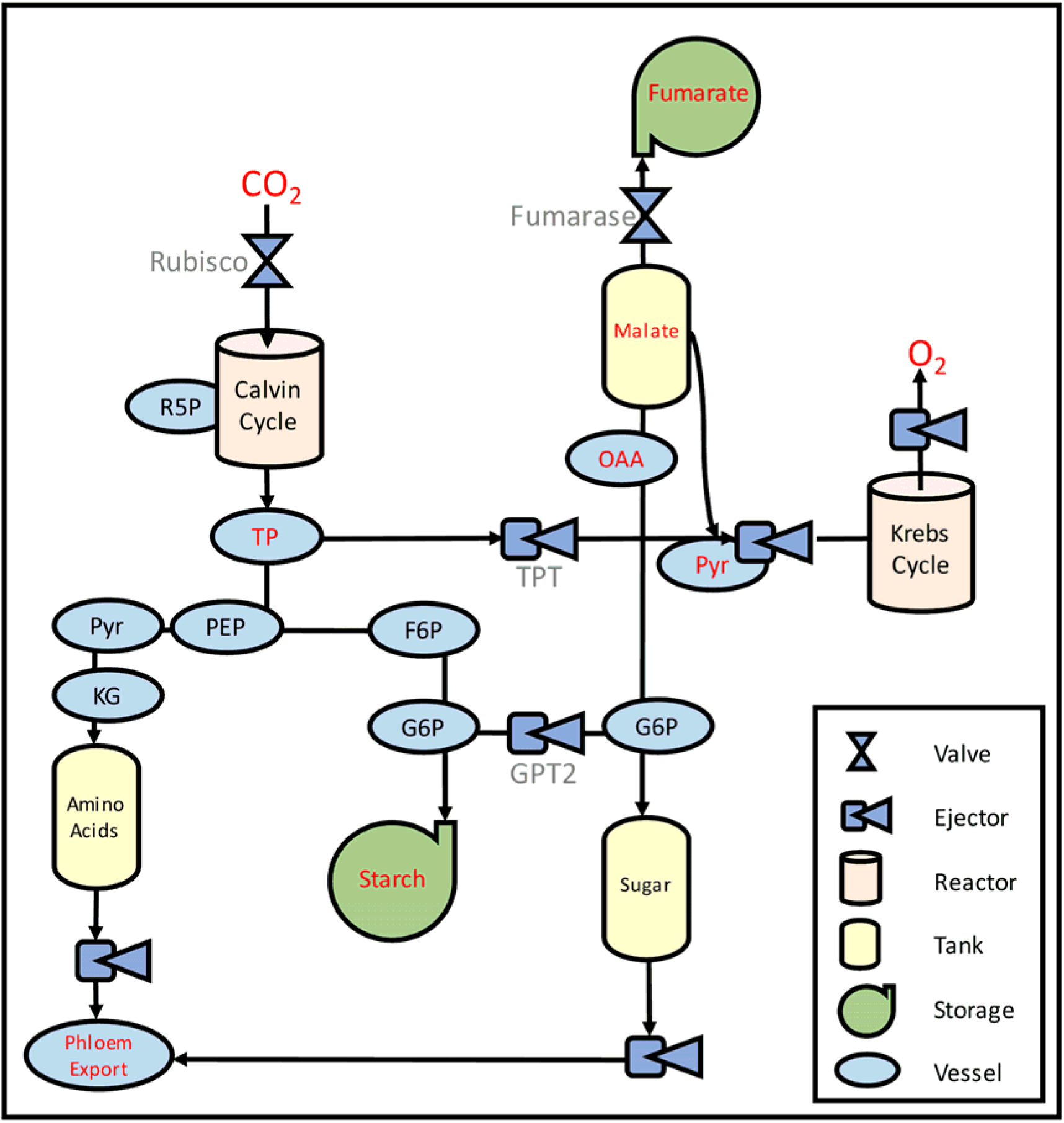
A simplified engineering system of plant primary metabolism connecting high-risk metabolites and their carbon sinks. All high-risk metabolites, as identified in Table 2 were included: R5P (ribose-5-phosphate), TP (glyceraldehyde 3-phosphate), PEP (phosphoenolpyruvate), Pyr (pyruvate), KG (α-ketoglutarate), F6P (fructose 6-phosphate), G6P (glucose 6-phosphate), OAA (Oxaloacetate) and Malate. Only branch and end point metabolites were included in the kinetic model and are shown in red.

Malate is a central metabolite in carbon metabolism, and alterations in its concentration levels could alter different cellular processes. Malate is not only an intrinsic part of the malate/OAA shuttle, participating in redox cellular regulation (reviewed by Selinski and Scheibe, 2018), but was also recently linked to membrane signalling under stress (Gilliham and Tyerman, 2016). In this model, the Aluminium-Activated Membrane Transporters (ALMT) are proposed to sense and signal metabolic status through the export of malate from the cytosol to the apoplasm (Ramesh et al. 2015). In particular, ALMT12 loss-of-function alters stomata dynamics (Meyer et al. 2010; Sasaki et al. 2010), with clear implications for photosynthesis and water stress responses.

To date, there is little direct experimental evidence for the presence of a cytosolic fumarase enzyme in plant species other than *Arabidopsis thaliana* (Chia et al. 2000). However, a recent phylogenetic study demonstrated that several close relatives of A. thaliana possess orthologues of the *fum2* gene (Zubimendi et al., 2018). This study also shows that other plant species obtained the gene through convergent evolution. Because cytosolic fumarase has recently evolved in Arabidopsis and its relatives (Zubimendi et al., 2018), it is likely that other mechanisms regulate the concentration of high-risk metabolites (like malate) in other species. For instance, phosphoenolpyruvate carboxylase (PEPC), which produces malate from oxaloacetate, is inhibited by malate across many plant species (O’Leary et al., 2011). This negative feedback loop could have the same effect as a fumarate sink, in that it maintains a constant accumulation of malate, even when the carbon influx is changing. This regulatory mechanism, however, is intricately dependent on phosphorylation and cellular pH (O’Leary et al., 2011), both of which may also be affected by environmental conditions. Thus, the evolution of a cytosolic fumarase fail-safe provides an alternative control mechanism regulating the malate concentration, while at the same time maintaining metabolic fluxes and allowing efficient storage of fixed carbon.

Although we have focused on malate as a high-risk metabolite, our analysis has identified other high-risk metabolic candidates. Interestingly, some of them have previously been noted as critical components of acclimation (Timm et al., 2012; Dyson et al., 2015; Weise et al., 2019). For example, it was shown that expression of GTP2, a chloroplast glucose 6-phosphate/phosphate translocator, is required for acclimation to high light in the Arabidopsis accession Wassilewskija-4 (Dyson et al., 2015). Our FMEA highlights glucose 6-phosphate in the chloroplast and in the cytosol as high-risk components, which are upstream of the starch and sucrose carbon sinks in the metabolic network (Fig. 7). GPT2 allows for appropriate distribution of carbon between these two sinks. When this control mechanism is broken, the two glucose 6-phosphate metabolite concentrations cannot be regulated and acclimation to high light is affected (Dyson et al., 2015). Other high-risk metabolites identified by our FMEA, such as glyceraldehyde 3-phosphate, pyruvate, phosphoenolpyruvate, and α-ketoglutarate, may also play important roles in maintaining system reliability under changing environmental conditions. Further research needs to be done to characterize the relevance of these metabolites, highlighted by our FMEA approach, with regards to stress acclimation responses in plants.

## Conclusion

Building on our previous research that cytosolic fumarate accumulation is required for photosynthetic acclimation to cold conditions in Arabidopsis plants; we have here demonstrated that cytosolic fumarate accumulation is also necessary for photosynthetic acclimation to warm. This finding excludes the flux to cytosolic fumarate and the concentration of fumarate in the cell as a direct temperature signal. Adapting a reliability engineering technique (FMEA), we instead propose FUM2 as a metabolic fail-safe that maintains system stability and allows for constant malate accumulation across changing environmental conditions. The implementation of FMEA in this study gives a theoretical framework to quantify the risk associated with metabolic alterations, which could also be the result of gene deletions or insertions. We believe that this novel approach holds the potential to be used in the identification of new metabolic signals participating in plants acclimation responses to a fluctuating environment.

## Acknowledgements

HAH is supported by a Biotechnology and Biological Sciences Research Council (BBSRC) Doctoral Training Partnership stipend (BB/M011208/1) and PIC is supported by BBSRC research grant (BB/S009078/1). We thank Armida-Irene Gjindali for proof-reading the final manuscript and for useful discussions.

## Author Contributions

HAH, JMS and GNJ conceived the study. HAH conducted the experiments and developed the FMEA framework. PIC contributed to the analysis of the data. All authors co-wrote the manuscript.

